# CLE42 Delays Leaf Senescence by Antagonizing Ethylene Pathway in *Arabidopsis*

**DOI:** 10.1101/2022.02.27.481379

**Authors:** Yi Zhang, Shuya Tan, Yuhan Gao, Chengcheng Kan, Hou-Ling Wang, Qi Yang, Xinli Xia, Takashi Ishida, Shinichiro Sawa, Hongwei Guo, Zhonghai Li

## Abstract

Leaf senescence is the final stage of leaf development and is influenced by numerous internal and environmental factors. CLE family peptides are plant-specific peptide hormones that regulate various developmental processes. However, the role of CLE in regulating leaf senescence remains unclear. Here, we found that CLE42 is a negative regulator of leaf senescence by using a CRISPR/Cas9-produced CLE mutant collection. The *cle42* mutant displayed earlier senescence phenotypes, while overexpression of *CLE42* delayed age-dependent and dark-induced leaf senescence. Moreover, application of the synthesized 12-aa peptide (CLE42p) also delayed leaf senescence under natural and dark conditions. CLE42 and CLE41/44 displayed functional redundancy in leaf senescence, and the *cle41 cle42 cle44* triple mutant displayed more pronounced earlier senescence phenotypes than any single mutant. Analysis of differentially expressed genes obtained by RNA-Seq methodology revealed that ethylene pathway was suppressed by overexpressing *CLE42*. Moreover, CLE42 suppressed ethylene biosynthesis and thus promoted the protein accumulation of EBF, which in turn decreased the function of EIN3. Accordingly, mutation of EIN3/EIL1 or overexpression of *EBF1* suppressed the earlier senescence phenotypes of the *cle42* mutant. Together, our results reveal that the CLE peptide hormone regulates leaf senescence by communicating with ethylene pathway.

## Introduction

Plant leaves capture energy from sunlight to fix carbon in glucose, which is the source of food for most organisms on earth (Hohmann-Marriott & Blankenship, 2011). As leaves age, their photosynthetic capacity gradually decreases, accompanied by the catabolism of macromolecules, including nucleic acids, proteins, and lipids. The nutrients released from senescent leaves are recycled to other developing organs, such as young leaves, flowers, and seeds, leading to an increase in reproductive success (Guo & Gan, 2005; Lim *et al.*, 2007). Efficient senescence is essential for maximizing viability in the next generation or season, whereas premature senescence caused by various internal or environmental signals reduces the yield or fresh product quality of agronomically important crop plants (Hortensteiner & Feller, 2002). Thus, the appropriate onset and progression of leaf senescence are critical for plant fitness (Uauy *et al.*, 2006).

Leaf senescence is not a passive process but a genetically controlled developmental process (Nam, 1997; Guo & Gan, 2005; Lim & Nam, 2005). In model plants such as Arabidopsis, tomato, and rice, a number of components in the regulation of leaf senescence were identified by forward genetic screenings (Lim *et al.*, 2007; Woo *et al.*, 2016; Li *et al.*, 2020). Through multi-omics studies such as genomics, transcriptomics, proteomics, and metabolomics, thousands of senescence-associated genes (SAGs) were identified in dozens of plant species (Woo *et al.*, 2013; Kim *et al.*, 2016). Reverse genetics studies, including the construction of gain- or loss-of-function mutants, have also identified hundreds of functional SAGs (Li *et al.*, 2020).

Although leaf senescence is mainly controlled by developmental age, it is also influenced by an array of internal and external signals (Guo & Gan, 2005; Lim *et al.*, 2007). Plant hormones are major players regulating various stages of leaf senescence, including the onset, progression, and terminal phase of senescence. Ethylene, jasmonic acid (JA), abscisic acid (ABA), salicylic acid (SA), and strigolactone promote leaf senescence, whereas cytokinin, auxin, and gibberellic acid (GA) delay this process (Gan & Amasino, 1995; Lim *et al.*, 2007; Woo *et al.*, 2019). Among them, ethylene plays an important regulatory role in leaf senescence and fruit ripening (Abeles *et al.*, 1988). Exogenous application of ethylene promotes leaf senescence, while spraying ethylene synthesis or signaling inhibitors delays senescence (Abeles *et al.*, 1988). Forward genetic screening revealed that EIN2 (ETHYLENE-INSENSITIVE2), the core member of the ethylene signaling pathway, is a positive regulator of leaf senescence (Oh *et al.*, 1997). Mutation of EIN2 delayed leaf senescence (Oh *et al.*, 1997; Kim *et al.*, 2009), while overexpression of *EIN2* promoted this process (Li *et al.*, 2014). EIN3, a master transcription factor in the ethylene pathway, is also a positive regulator of leaf senescence (Li *et al.*, 2013). EIN3 directly binds to the promoter of *miR164* and inhibits its expression, thereby indirectly promoting the expression of *ORE1*, a core component in the regulation of leaf senescence (Li *et al.*, 2013; Kim *et al.*, 2014). In addition, EIN3 directly regulates the expression of chlorophyll catabolic genes and promotes leaf senescence (Qiu *et al.*, 2015). Ethylene also interacts with JA, SA, and strigolactone signaling pathways to coordinate the leaf senescence process (Li *et al.*, 2013; Ueda & Kusaba, 2015; Wang *et al.*, 2021; Yu *et al.*, 2021).

Plant polypeptide hormones are a class of small molecule signaling substances that play an important role in regulating plant growth and development and the response to environmental stresses (Fiers *et al.*, 2007; Betsuyaku *et al.*, 2011; Hirakawa & Sawa, 2019). Since the discovery of the small peptide hormone systemin in tomato (Pearce *et al.*, 1991), an increasing number of plant peptide signaling molecules have been identified (Hirakawa & Sawa, 2019). The CLAVATA3 (CLV3)/ENDOSPERM SURROUNDING REGION (ESR) (CLE) peptide family is one of the largest families of plant peptide hormones (Fletcher, 2020). The CLE gene encodes a precursor protein with a full length of approximately 100 amino acids (aa), which is processed by an unknown protease to form a mature peptide with a length of 12 to 14 aa (Ito *et al.*, 2006; Fletcher, 2020). CLE mature peptides bind to different membrane-bound receptors and thus transduce signals into the cell to regulate plant development by modulating downstream transcription factors or communicating with canonical plant hormones (Etchells *et al.*, 2012; Gao & Guo, 2012; Zhang *et al.*, 2016; Mou *et al.*, 2017; Fletcher, 2020). CLE family members are expressed in multiple tissues and are involved in regulating various aspects of plant development (Fiers *et al.*, 2007; Yamaguchi *et al.*, 2016), but their role in leaf senescence remains largely unclear.

A major problem in dissecting the functions of peptide hormones is the lack of loss-of-function mutants generated by canonical mutagenesis methods due to their small gene size (Yamaguchi *et al.*, 2017). Fortunately, CRISPR/Cas9-mediated gene editing technology overcomes this difficulty and has been used to generate loss-of-function mutants of CLE family genes (Yamaguchi *et al.*, 2017). Using a CRISPR/Cas9-produced CLE mutant collection, we found that the leaves of CLE42 loss-of-function mutants senesced prematurely, whereas overexpression of *CLE42* significantly delayed the leaf senescence process, suggesting that CLE42 is a negative regulator of senescence. Transcriptome and genetic analyses revealed that CLE42 delays leaf senescence by suppressing the ethylene pathway.

## Materials and Methods

### Plant materials and growth conditions

#### Arabidopsis thaliana

Columbia-0 (Col-0) was used as the wild type in this study. The CRISPR/Cas9-produced CLE mutant collection was described previously (Yamaguchi *et al.*, 2017). The plant materials *pxy-3* and *pxy pxl1 pxl2* (Smit *et al.*, 2020), *ein3-1 eil1-1* (Alonso *et al.*, 2003), and *ACS octuple* mutant (*acs2-1 acs4-1 acs5-2 acs6-1 acs7-1 acs9-1 amiRacs8 acs11*) (Tsuchisaka et al., 2009) used in this study were described previously. Seeds were surface-sterilized and plated on Murashige and Skoog (MS) medium (4.3 g/L MS salts, 1% sucrose [pH 5.7 to 5.8], and 8 g/L agar). The plates were then stored for 2 d at 4 °C before exposing them to white light (PAR 100 to 150 μE m^−2^ s^−2^). After 3-4 d, light-grown seedlings were transferred to soil and grown at 22 °C under a 16 h light/8 h dark cycle (PAR 100 to 150 μE m^−2^ s^−2^). For the triple response assay, genotypes were grown on MS medium supplied with or without the indicated concentration of 1-aminocyclopropane-1-carboxylic acid (ACC) (Sigma-Aldrich, CAS 22059-21-8).

### Plasmid construction and transformation

To generate *35S:GFP-CLE42*/Col-0, the full-length *CLE42* cDNA sequence was amplified and then inserted into the pEGAD vector (Cutler *et al.*, 2000). To construct the β-estradiol inducible *CLE42* overexpression lines (*iCLE42*), the CLE42 (linked with 3×Flag) CDS was amplified and then introduced into the pER8 vector (Zuo *et al.*, 2000). Amplified fragments were inserted into the respective vectors by using the In-Fusion enzyme (Takara Bio, USA). For construction of CRISPR/cas9-mediated *CLE42* knockout plants, a target site of *CLE42* designed using CRISPR-P (http://crispr.hzau.edu.cn/CRISPR2/) was used (Table S1), and the homozygous plants were identified by sequencing. All constructs were transformed into *Agrobacterium tumefaciens* cells (strain GV3101), which were then used to transform Col-0 plants using the floral dip method (Clough & Bent, 1998). All primer sequences used here are listed in Table S1.

### Leaf senescence assays under natural and stress conditions

All senescence experiments were performed using the third and fourth rosette leaves. Age-dependent leaf senescence was evaluated as described previously (Woo *et al.*, 2001). Leaves of individual plants at the indicated developmental stages were detached, photographed, and used to assess the chlorophyll content, the photochemical efficiency of PSII (Fv/Fm) and *SAG* expression. For the hormone-, salt-, or dark-induced leaf senescence assay, leaves at 12 d after emergence (DAE) were detached and floated on 5 mM MES buffer (pH 5.7) in the presence of hormone or NaCl at 22 °C in the dark (Woo *et al.*, 2004).

### RNA extraction and quantitative Real-Time PCR analysis

Total RNA was extracted using plant RNA kits (ER301; TransGen Biotech, China), and cDNA was synthesized using a cDNA synthesis kit (AT341; TransGen Biotech, China). Quantitative RT-PCR (qPCR) analysis was performed to determine gene expression levels using an ABI7500 Real-Time PCR system with a qPCR kit (W812014, Welab Biotech, China), and then relative mRNA quantities were calculated from the average values using the ΔΔC_T_ method (Schmittgen & Livak, 2008). All primers were shown in Table S1.

### RNA-sequencing analysis

The third and fourth rosette leaves of 24-d-old Col-0 and *CLE42ox* plants were collected and ground into a powder in liquid nitrogen. All samples were collected in three biological replicates. Total RNA was extracted using a RNeasy Plant Kit (Qiagen, cat. nos. 74904), and the quality and quantity of RNA were checked using an Agilent 2100 Bioanalyzer (Agilent Technologies, Inc., Santa Clara, CA, USA). RNA-Seq data were generated with an Illumina HiSeq™ 2500 sequencing platform (Biomarker Ltd., Beijing, China). Raw reads (fastq format) were trimmed and filtered using in-house Perl scripts (Biomarker Ltd, China). The reads were then mapped to the Arabidopsis reference genome using the Hisat2 algorithm. DEGs were filtered using the following criteria: |Log2 (fold change)| > 1.0, P < 0.05. Gene Ontology (GO) enrichment analysis was performed by using the GO database (http://geneontology.org/). Default parameters were used for all bioinformatics software.

### Histochemical analysis of GUS activity

GUS staining was performed as previously described (Jefferson *et al.*, 1987). We incubated tissues or seedlings with GUS staining solution (100 mM Na3PO4 [pH 7.0], 1 mM potassium ferricyanide, 1 mM EDTA, 1% Triton X-100, 1 mM potassium ferrocyanide, and 1 mg/mL 5-bromo-4-chloro-3-indolyl-β-D-glucuronide) overnight at 37 °C and then incubated them for hours in ethanol to eliminate chlorophyll before photographing. GUS quantification was carried out using 4-methylumbelliferyl-β-D-glucuronide (4-MUG; Sigma-Aldrich) as the substrate.

### Western blotting

One hundred milligrams of tissues or seedlings was ground into a powder in liquid nitrogen, and 100 μL of 2× protein loading buffer was then added and boiled for 5 min. Subsequently, 10 μL per sample was analyzed by western blotting using monoclonal anti-GFP antibodies conjugated to horseradish peroxidase (HRP; Sigma-Aldrich) and separated by 8% SDS-PAGE. Coomassie brilliant blue (CBB) was the total protein loading control.

### Measurement of ethylene production

The third/fourth rosette leaves of Arabidopsis were used for quantification of ethylene production. The detached leaves treatment with or without CLE42p solution were weighed and incubated in a vial at 22°C under a 16 h light/8 h dark cycle. Ethylene production after 24 h of incubation was measured with gas chromatography as previously described (Tsuchisaka et al., 2009; Qiu et al., 2015).

### Chromatin immunoprecipitation (ChIP)-qPCR

Two grams of 5-d-old *35S:EIN3-GFP* seedlings treated with CLE44p or ACC were used for ChIP experiments as previously described (Saleh *et al.*, 2008). GFP antibody was used to immunoprecipitate the protein-DNA complex. The enrichment of DNA fragments was determined by qPCR. All primers used here are listed in Table S1.

## Results

### CLE42 negatively regulates age-dependent and dark-induced leaf senescence

To investigate the function of CLE family peptides in leaf senescence, we first performed a senescence phenotypic screening by using a CRISPR/Cas9-produced CLE mutant collection (Yamaguchi *et al.*, 2017). The mutants were grown in soil under long-day conditions (16 h light/8 h dark) alongside wild-type Col-0. We found that the *cle42* mutant displayed a premature senescence phenotype (Fig. 1a), suggesting that CLE42 is a negative regulator of leaf senescence. The CLE42 peptide and TRACHEARY ELEMENT DIFFERENTIATION INHIBITION FACTOR (TDIF; CLE41/CLE44 peptide) act as inhibitors of tracheary element differentiation (Yaginuma *et al.*, 2011; Etchells *et al*., 2016). To further determine its function, we generated transgenic plants overexpressing *CLE42* fused with GFP driven by the *35S* promoter (*35S:GFP-CLE42*, *CLE42ox*), and two transgenic lines (*CLE42ox* #1 and #2) were chosen for further analysis (Fig. S1). Remarkably, the leaves of *CLE42ox* plants displayed delayed senescence compared to those of Col-0 and *cle42* (Fig. 1a,b). We next examined in detail the senescence phenotypes of the fourth leaves of Col-0, *cle42*, and *CLE42ox* during their lifespans. The leaves of *cle42* mutants started to turn yellow from the tip at Day 22 (i.e., 22 d after emergence of the fourth leaves), whereas Col-0 and *CLE42ox* leaves turned yellow at Days 26 and 34, respectively (Fig. 1c), with lower chlorophyll contents (Fig. 1d) and photochemical efficiency of PSII (Fv/Fm) (Fig. 1e). Transcript levels of *SAG12*, a marker gene of leaf senescence, in the *cle42* mutant were more dramatically induced than those in Col-0 and *CLE42ox* (Fig. 1f). Additionally, the transcripts of *CLE42* and *CLE41/44* decreased with leaf age (Fig. S2), raising the possibility that they act as suppressors of leaf senescence.

**Fig. 1.**
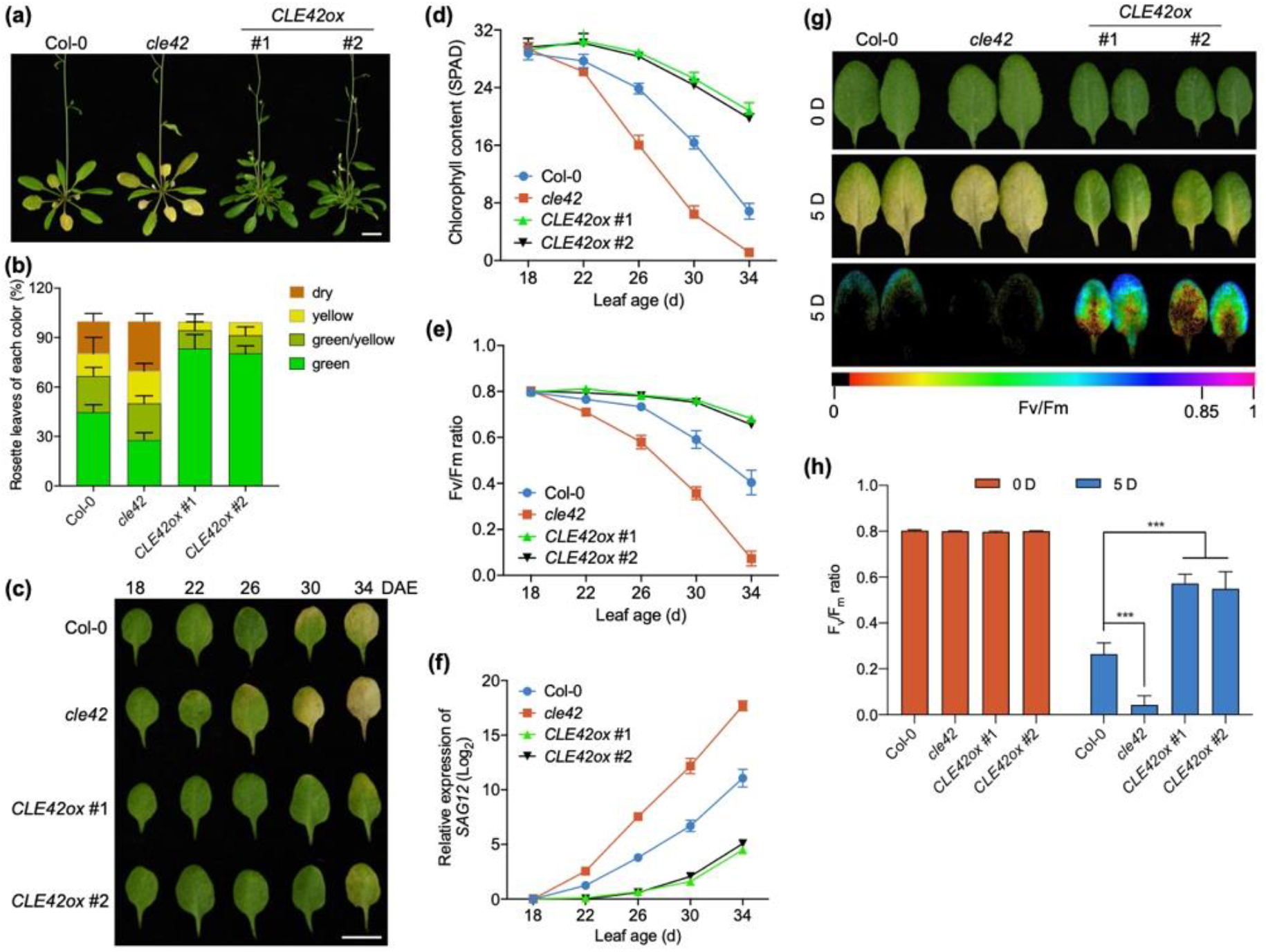
CLE42 negatively regulates age-dependent and dark-induced leaf senescence. (a) The senescence phenotypes of 40-d-old wild-type Col-0, *cle42*, and *CLE42ox*. Two independent lines, *CLE42ox* #1 and #2, were examined. Bar, 1 cm. (b) quantification of the leaf senescence phenotypes of the third/fourth rosette leaves shown in (a). (c) Phenotypes of the third/fourth rosette leaves of Col-0, *cle42*, and *CLE42ox* at the indicated age. Representative leaves are shown. Bar, 1 cm. (d-f) Chlorophyll content (d), Fv/Fm ratio (e), and *SAG12* expression (f) in the leaves shown in (c). The bars indicate the mean ± SD (n = 20). (g) The senescence phenotypes and Fv/Fm ratio of Col-0, *cle42*, and *CLE42ox* leaves upon dark treatment for 4 days. (h) Fv/Fm ratio of leaves shown in (g). The bars indicate the mean ± SD (n = 20). ***P < 0.001, two-way ANOVA.

To further assess the role of CLE42 in leaf senescence, we examined the senescence phenotypes of *cle42* and *CLE42ox* under dark conditions. The third rosette leaves detached from 21-d-old Col-0, *cle42*, and *CLE42ox* plants were placed in the dark. We found that *cle42* exhibited significantly faster dark-induced senescence, while *CLE42ox* showed significantly delayed dark-induced senescence than Col-0 (Fig. 1g,h). We also tested the effects of other senescence-regulating signals, such as plant hormones, salt stress, and H_2_O_2_, on the senescence phenotypes of *cle42* and *CLE42ox*. Compared with the Col-0 and *cle42* mutants, the leaves of *CLE42ox* displayed delayed senescence phenotypes upon treatment with ABA, JA, ACC (ethylene precursor), NaCl, or H_2_O_2_ under dark conditions (Fig. S3). Together, these results indicate that CLE42 delays both age-dependent and dark- and stress-induced leaf senescence.

### CLE41/44 and PXY are involved in leaf senescence

The mature form of CLE42 is a dodecapeptide (HGVPSGPNPISN) and differs from CLE41/CLE44 (HEVPSGPNPISN) only in the second position. Both forms play an essential role in regulating cell division in vascular meristems by binding to the leucine-rich repeat receptor kinase (LRR-RK) PXY (PHLOEM INTERCALATED WITH XYLEM) (Fisher & Turner, 2007; Hirakawa *et al.*, 2008; Zhang *et al.*, 2016). To examine whether there is genetic redundancy between CLE42 and CLE41/CLE44 in regulating leaf senescence, we generated *cle41 cle42 cle44* triple mutants by crossing (Fig. S4). The onset of leaf senescence started earlier in *cle41 cle42 cle44* triple mutants than in any single mutant (Fig. 2a), with a pronounced decrease in the chlorophyll content and Fv/Fm as well as an earlier induction of *SAG12* expression (Fig. 2b–d), indicative of the existence of genetic redundancy. Interestingly, the *cle42* mutant displayed stronger senescence phenotypes than *cle41* and *cle44*, with the lowest chlorophyll contents and Fv/Fm (Fig. 2b,c), suggesting that CLE42 plays a dominant role in leaf senescence. As expected, mutation of *PXY* accelerated leaf senescence (Fig. 2a), suggesting that the CLE41/42/44-PXY module prevents leaf from premature senescence in Arabidopsis.

**Fig. 2.**
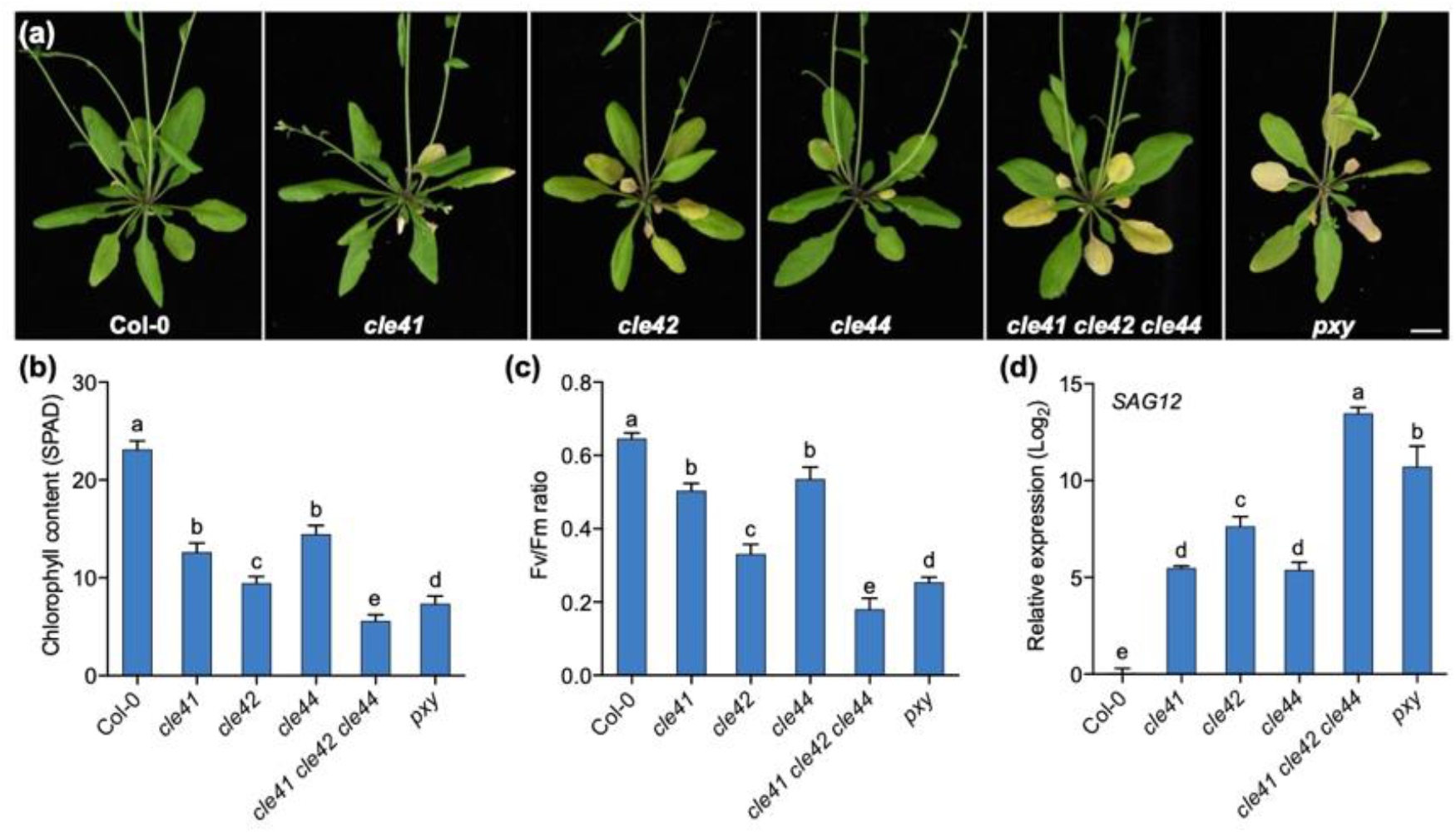
CLE41/42/44-PXY module is involved in leaf senescence. (a) The senescence phenotypes of 35-d-old Col-0, *cle41*, *cle42*, *cle44*, *cle41 cle42 cle44*, and *pxy*. Bar, 1 cm. (b-d) Chlorophyll content (b), Fv/Fm ratio (c), and *SAG12* expression (d) in the fourth leaves shown in (a). Different letters above the bars indicate statistically significant differences (adjusted P < 0.05, one-way ANOVA).

In Arabidopsis, two kinases, PXY-like 1 (PXL1) and PXL2, are closely related to PXY (Fisher & Turner, 2007), and the CLE41/42/44 peptides also interact with PXL1/2 (Zhang *et al.*, 2016; Mou *et al.*, 2017). Single mutants of *pxl1* and *pxl2* did not exhibit obvious phenotypic changes in the vascular stem; however, the triple mutant *pxy pxl1 pxl2* showed enhanced alterations of the vascular phenotype compared with that of *pxy* (Etchells *et al.*, 2013), indicating that PXL1 and PXL2 function redundantly with PXY in the regulation of vascular tissue development. Unexpectedly, no evidently accelerated senescence phenotype was observed in *pxy pxl1 pxl2* leaves in comparison with *pxy* leaves (Fig. S5), indicating that PXL1 and PXL2 play a minor role in regulating leaf senescence. Interestingly, the leaves of *cle41 cle42 cle44* showed stronger earlier senescence symptoms than those of *pxy* or *pxy pxl1 pxl2*, with lower chlorophyll contents and Fv/Fm, as well as earlier induction of *SAG12* expression (Fig. 2b–d; Fig. S5), suggesting that CLE41/CLE42/CLE44 regulates leaf senescence in a PXY-dependent and PXL-independent manner.

### CLE42p delays age-dependent and dark-induced leaf senescence

To determine whether the mature form of the CLE42 peptide is enough to function as a regulator of leaf senescence, we synthesized a 12-aa peptide (CLE42p, HGVPSGPNPISN) derived from the CLE domain of the CLEL42 precursor. Wild-type Col-0 and *cle42* plants were sprayed with water (Mock) or 5 μM CLE42p every 4 days. Remarkably, exogenous application of CLE42p significantly delayed leaf senescence in Col-0 plants and compensated for earlier senescence of *cle42* mutants (Fig. 3a). Time-course analyses revealed that treatment with CLE42p evidently delayed the leaf senescence process in Col-0 and *cle42* backgrounds, with higher levels of chlorophyll contents and Fv/Fm compared with the controls (Fig. 3b,c). Interestingly, the *cle42* mutant displayed senescence phenotypes similar to those of Col-0 upon treatment with CLE42p (Fig. 3a–c), implying that in vitro synthesized CLE42p is biologically functional and rescues the phenotypes of *cle42*. In contrast, the *pxy* mutant exhibited earlier leaf senescence phenotypes, with lower chlorophyll content and Fv/Fm, which was slightly rescued by CLE42p, implying that CLE42 regulates leaf senescence mainly *via* the receptor PXY. Inducible overexpression of *CLE42* delayed leaf senescence in Col-0 but not in the *pxy* background, thus supporting our hypothesis (Fig. 3d–f).

**Fig. 3.**
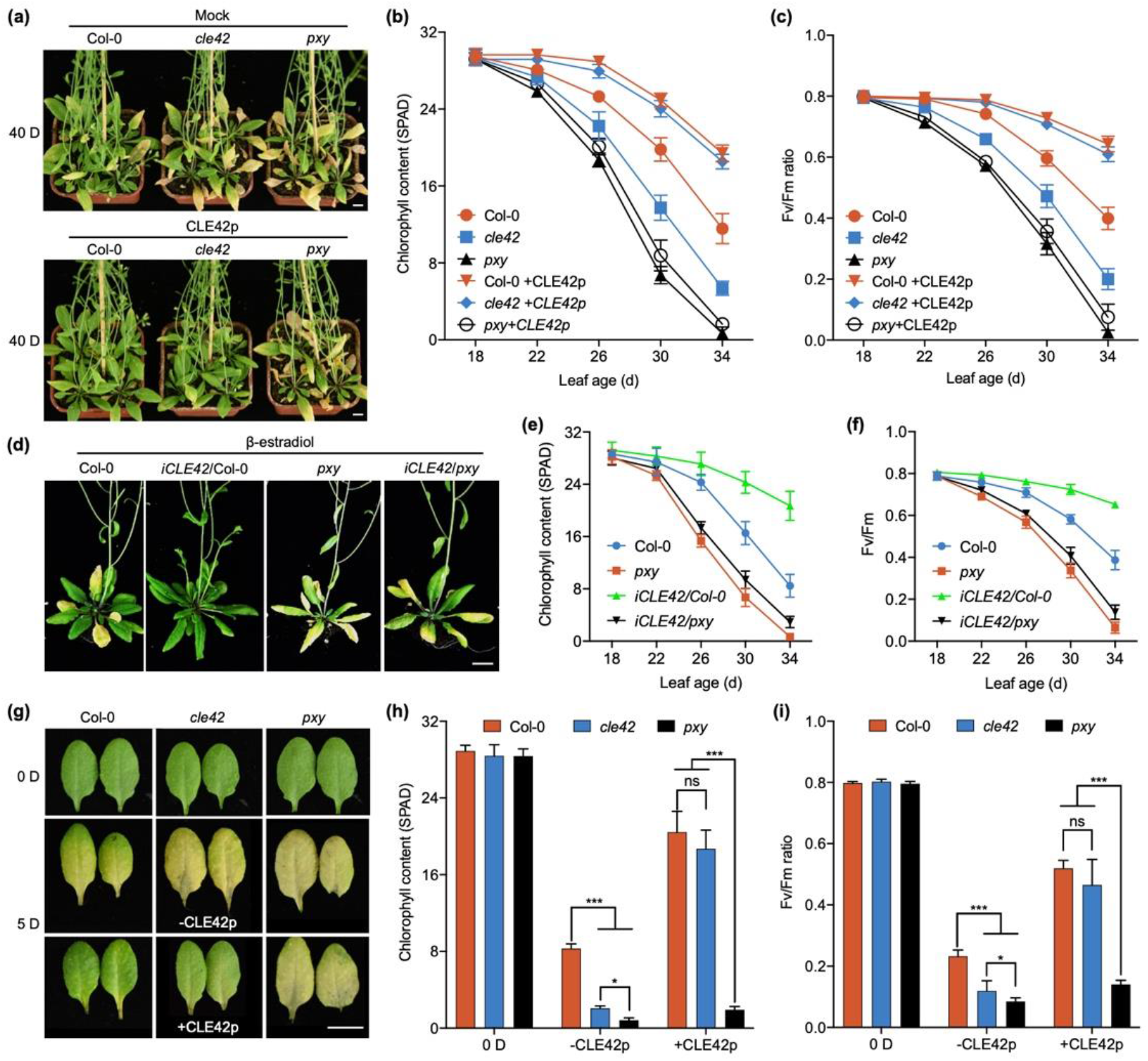
Exogenous application of CLE42p delays age-dependent and dark-induced leaf senescence. (a) The senescence phenotypes of 40-d-old wild-type Col-0 and *cle42* upon treatment with CLE42p. Col-0 and *cle42* were germinated on MS containing water (Mock) or 10 μM CLE42p, and the 5-day-old seedlings were transferred to soil and sprayed with CLE42p every 4 d. Bars, 1 cm. (b) and (c) Fv/Fm ratio (b) and chlorophyll content (c) in the leaves shown in (a) at the indicated age. The bars indicate the mean ± SD (n = 20). (d) The senescence phenotypes of 40-d-old Col-0, *iCLE42*/Col-0, *pxy*, and *iCLE42*/*pxy* plants upon treatment with β-estradiol. Seeds were germinated on MS medium containing 10 μM β-estradiol, and 5-d-old seedlings were transferred to soil and then sprayed with 10 μM β-estradiol every 4 days. Bar, 1 cm. (e) and (f) Chlorophyll content (e) and Fv/Fm ratio (f) in the leaves of Col-0, *iCLE42*/Col-0, *pxy*, and *iCLE42*/*pxy* plants at the indicated age. (g) The senescence phenotypes and Fv/Fm ratio of Col-0 and *cle42* leaves upon treatment with CLE42p under darkness for 4 days. Bars, 1 cm. (h) and (i) chlorophyll content (h) and Fv/Fm ratio (i) in the leaves shown in (g). The bars indicate the mean ± SD (n = 20). *P < 0.05, ***P < 0.001, two-way ANOVA.

We also examined the effect of CLE42p on leaf senescence under dark conditions. Compared with the controls, CLE42p evidently delayed the dark-induced senescence process in Col-0 and *cle42* backgrounds (Fig. 3g), with high levels of chlorophyll content and Fv/Fm (Fig. 3h,i), while slight influence was observed in *pxy* leaves. Similarly, synthetic CLE41/44p (HEVPSGPNPISN) was also able to restore the phenotypes of the *cle42* mutant under dark conditions (Fig. S6), which further suggests that there is functional redundancy between CLE42 and CLE41/44. Together, our data demonstrated that exogenous application of synthetic CLE42p is sufficient to delay both age-dependent and dark-induced leaf senescence.

### Transcriptome analysis reveals the CLE42-mediated signaling pathway

To reveal the molecular basis of CLE42-regulated leaf senescence, we performed transcriptome analysis of 21-day-old leaves of Col-0 and *CLE42ox* plants. Compared with Col-0, 2,495 differentially expressed genes (DEGs) were identified in *CLE42ox* leaves (|log2(FC)|>1, FDR<0.05) (Dataset S1). Gene Ontology (GO) enrichment analysis of the DEGs showed that the upregulated genes were highly enriched in the regulation of meristem growth and cell size, cell wall biogenesis, auxin polar transport, and xylem development (Fig. S7a), which is consistent with previous findings that TDIFs are involved in regulating meristem and xylem development. The downregulated genes were related to defense responses and hormone signaling pathways, such as JA, SA, and ethylene (Fig. S7b), suggesting that CLE42 delays leaf senescence by suppressing hormone signaling. In agreement with the delayed senescence phenotype of *CLE42ox*, many *SAG*s were downregulated, such as *SAG12*, *SAG13*, *ORE1*, *WRKY75*, *WRKY53*, and *YLS9* (Fig. S8).

A previous study showed that the TDIF-PXY signaling pathway may interact and suppress ET signaling in vascular development, as many ET-responsive factors were upregulated in the *pxy* mutant by microarray expression analysis (Etchells *et al.*, 2012). The triple response phenotype is commonly used as an ethylene-specific growth response in Arabidopsis and refers to the exaggerated apical hooks, shortened hypocotyls, and roots of dark-grown seedlings exposed to ethylene or treated with the ethylene precursor ACC (Bleecker *et al.*, 1988). Overexpression of *CLE42* conferred significant attenuation of triple response phenotypes, with longer hypocotyls, compared with wild-type seedlings upon treatment with low concentrations of ACC (0.1-5 μM) (Fig. S9). In contrast, seedlings of *cle42* mutants displayed an enhanced response to low concentrations of ACC (0.1-5 μM) (Fig. S9), suggesting that CLE42 is involved in the ethylene signaling pathway. Notably, although seedlings of *cle41*, *cle42* and *cle44* single mutants showed enhanced sensitivity to ethylene compared with Col-0, the triple response of *cle42* seedlings was the most obvious (Fig. S10), implying that CLE42 may act as a prominent player in the communication between TDIF-PXY and the ethylene signaling pathway.

### CLE42 decreases the function of EIN3 by promoting the protein accumulation of EBF

Because EIN3 is a master transcription factor of the ethylene signaling pathway and a positive regulator of leaf senescence, we next investigated the effect of CLE42p treatment on its function. To this end, two transgenic lines that harbor the GUS reporter gene driven by five tandem repeats of the EIN3 binding site (EBS) followed by the minimal 35S promoter, *5×EBS:GUS*/Col-0 and *5×EBS:GUS/EIN3ox* (Stepanova *et al.*, 2007), were used to examine the function of EIN3 in the absence or presence of CLE42p. As expected, a high level of EIN3 function was observed in the seedlings of *5×EBS:GUS/EIN3ox* in comparison with *5×EBS:GUS*/Col-0 (Fig. 4a,b). Interestingly, the functions of EIN3 were significantly reduced in the seedlings of *5×EBS:GUS*/Col-0 and *5×EBS:GUS/EIN3ox* upon treatment with CLE42p compared with the controls (Fig. 4a), which was further supported by the quantification assays of GUS activity (Fig. 4b). These data suggest that CLE42 suppresses the function of EIN3.

**Fig. 4.**
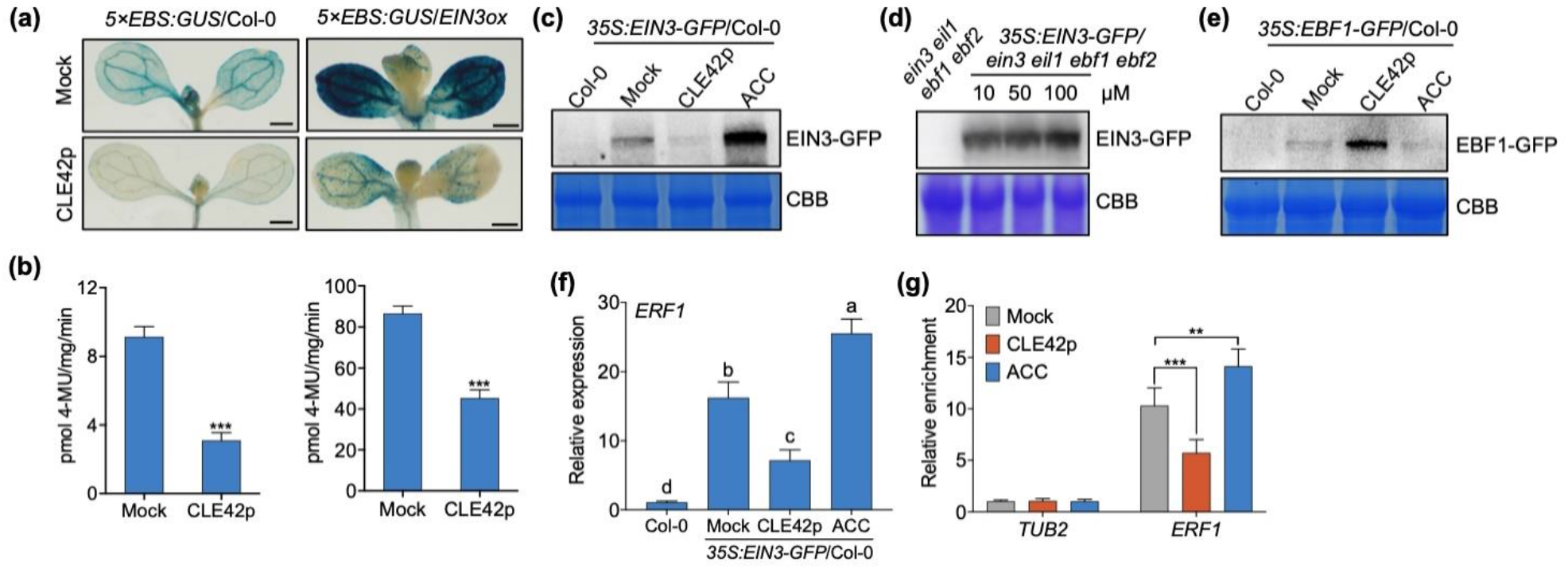
CLE42 represses the function of EIN3. (a) Histochemical analysis of *5×EBS:GUS*/Col-0 and *5×EBS:GUS*/*EIN3ox* transgenic plants upon CLE42p treatment. Bars, 500 μm. (b) GUS activity in seedlings shown in (A) was calculated as picomoles of 4-MU per mg protein per min. ***P < 0.001, one-way ANOVA. (c) CLE42p reduces EIN3 protein accumulation in the wild type. Five-day-old seedlings were treated with 10 μM CLE42p or ACC for 24 h. Protein was extracted and subjected to immunoblots using anti-GFP antibody. Coomassie brilliant blue (CBB) staining was used as the protein loading control. (d) CLE42p-induced EIN3 protein degradation depends on EBF1/2. *35S:EIN3-GFP/ein3 eil1 ebf1 ebf2* seedlings treated with 10, 50 and 100 μM CLE42p for 24 h were used for protein extraction. (e) CLE42p promotes EBF1 protein accumulation. Five-day-old seedlings were treated with 10 μM CLE42p or ACC for 24 h. Protein was extracted and subjected to immunoblots using anti-GFP antibody. Coomassie brilliant blue (CBB) staining was used as the protein loading control. (f) The transcript level of *ERF1* in Col-0 and *35S:EIN3-GFP* (*EIN3ox*) upon 10 μM CLE42 or ACC treatment. The bars represent the standard deviation of three technical replicates. Different letters above the bars indicate statistically significant differences (adjusted P < 0.05, one-way ANOVA). (g) ChIP-qPCR was performed to examine the relative EIN3 binding to the *ERF1* promoter under CLE42p treatment. An anti-GFP monoclonal antibody was used for DNA immunoprecipitation from 5-d-old *EIN3ox* transgenic plants. The relative enrichment of EIN3 binding to the *ERF1* promoter was normalized to *TUBULIN2* (*TUB2*). The bars represent the standard deviation of three technical replicates. Each experiment was repeated at least three times with similar results. **P < 0.01, ***P < 0.001, one-way ANOVA.

We next investigated how EIN3 function is modulated by CLE42p. To this end, we monitored the level of EIN3 protein using *35S:EIN3-GFP*/Col-0 plants with anti-GFP antibody in response to CLE42p treatment. In agreement with previous findings, the level of EIN3 protein dramatically increased upon treatment with 1-aminocyclopropane-1-carboxylic acid (ACC), a key precursor of ethylene biosynthesis. In contrast, the protein accumulation of EIN3 was evidently decreased in the presence of CLE42p (Fig. 4c), suggesting that the decreased transcriptional functioning of EIN3 was caused by the decreased EIN3 protein level. We also checked the levels of EIN3 mRNA, and no obvious change was observed in Col-0 seedlings upon treatment with CLE42p after 6 h (Fig. S11), suggesting that CLE42p regulates EIN3 function at the protein level. However, no change was observed in the protein levels of EIN3 in the transgenic plants *35S:EIN3-GFP*/*ein3 eil1 ebf1 ebf2* upon treatment with CLE42p (Fig. 4d). EIN3-binding F-box 1 (EBF1) and EBF2 are key components of the ethylene signaling pathway, with distinct but overlapping roles in mediating the degradation of EIN3 protein through the Ub/proteasome pathway (Guo & Ecker, 2003; Potuschak *et al.*, 2003; Gagne *et al.*, 2004). These results revealed that EBF1/2 are required for the CLE42p-induced degradation of EIN3 protein. Next, we examined the levels of EIN3-binding F-box 1 (EBF1) protein after CLE42p treatment using *35S:EBF1-GFP*/Col-0 plants with anti-GFP antibody. Western blot assays showed that the protein levels of EBF1 remarkably increased upon CLE42p treatment (Fig. 4e), suggesting that CLE42p promotes the degradation of EIN3 protein by inducing the accumulation of EBF proteins.

To further investigate the effect of CLE42 on EIN3 function, we performed qPCR analysis to examine the expression of *ETHYLENE RESPONSE FACTOR1* (*ERF1*), a well-known downstream target of EIN3 and a positive regulator of ethylene signaling (Solano *et al.*, 1998). Treatment with ACC increased the transcription of ERF1, which was significantly decreased by CLE42p (Fig. 4f). Interestingly, the expression levels of *ERF1*, *ERF59/ORA59* and *PDF1.2* were significantly downregulated in *CLE42ox* plants compared with Col-0 plants (Fig. S12), suggesting that endogenous CLE42 suppresses the activity of EIN3. We next performed chromatin immunoprecipitation (ChIP) analysis to determine the effect of CLE42 on the binding activity of EIN3 to *ERF1* using *35S:EIN3-GFP*/Col-0 plants. As expected, EIN3 could directly bind to the promoter of ERF1, which was enhanced by ACC treatment (Fig. 4g). In contrast, CLE42p treatment evidently compromised the binding activity of EIN3 to *ERF1* (Fig. 4g), suggesting that CLE42p suppresses the protein stability of EIN3 and in turn affects its activity. Together, these results suggest that CLE42 suppresses the stabilization and function of EIN3 by promoting the protein accumulation of EIN3-targeting F-box proteins.

The above data pushed us to investigate how CEL42 affects the protein stability of EBF1. Interestingly, we found that the transcript level of *ACS2* was evidently suppressed in the leaves of *CLE42ox* compared with Col-0 (Fig. S13a). Moreover, ethylene production was evidently decreased in *CLE42ox* plants (Fig. S13b). In supporting these findings, exogenous application of CLE42p also suppressed the expressions of *ACS2* and the production of ethylene compared with the untreated plants (Fig. S13c,d). These results suggest that CLE42 promotes EBF1 protein accumulation possibly by suppressing endogenous ethylene production. Future studies are needed to investigate how CLE42 affects the expression of *ACS*s.

### Loss-of-function of EIN3/EIL1 or overexpression of *EBF1* suppresses the earlier senescence phenotypes of the *cle42* mutant

To further explore the genetic regulatory relationship between CLE42 and EIN3/EIL1 in leaf senescence, we generated *cle42 ein3 eil1* plants by crossing the *cle42* plant with the *ein3 eil1* mutant, which was confirmed by the presence of triple response phenotypes and PCR sequencing (Table S1). Under long-day conditions, compared to Col-0, *cle42 ein3 eil1* plants showed an obvious delayed senescence phenotype similar to *ein3 eil1* mutants, indicating that the accelerated leaf senescence phenotype of *cle42* mutants was effectively suppressed by the mutation of EIN3/EIL1 (Fig. 5a), which was confirmed by age-dependent changes in the chlorophyll content and Fv/Fm (Fig. 5b,c). Next, a dark-induced leaf senescence assay was performed using Col-0, *cle42*, *ein3 eil1*, and *cle42 ein3 eil1* plants. Remarkably, *ein3 eil1* strongly suppressed the earlier senescence phenotype of *cle42* under darkness (Fig. 5d). The leaves of *ein3 eil1* and *cle42 ein3 eil1* showed higher levels of chlorophyll content and Fv/Fm than Col-0 and *cle42* leaves (Fig. 5e,f). These results suggest that mutation of EIN3 and EIL1 suppresses the earlier senescence phenotypes of *cle42* plants.

**Fig. 5.**
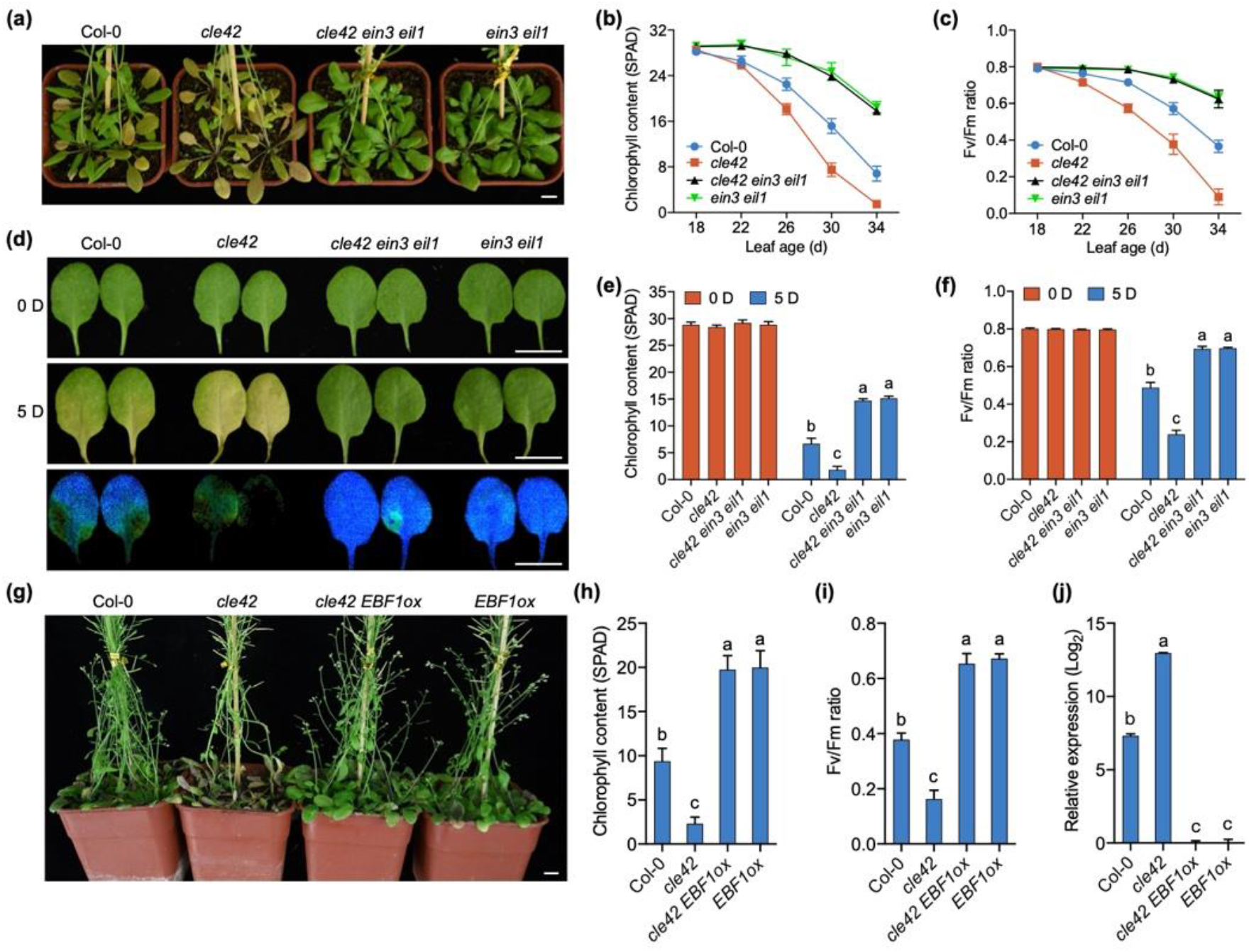
Mutation of EIN3/EIL1 or overexpression of *EBF1* suppresses the earlier senescence phenotypes of the *cle42* mutant. (a) The senescence phenotypes of 40-d-old wild-type Col-0, *cle42*, *ein3 eil1,* and *cle42 ein3 eil1*. Bar, 1 cm. (b) and (c) chlorophyll content (b) and Fv/Fm ratio (c) in the leaves shown in (a) at the indicated age. The bars indicate the mean ± SD (n = 20). (d) The senescence phenotypes and Fv/Fm ratio of Col-0, *cle42*, *ein3 eil1,* and *cle42 ein3 eil1* leaves upon dark treatment for 4 days. (e) and (f) chlorophyll content (e) and Fv/Fm ratio (f) in the leaves shown in (d). (g) The senescence phenotypes of 45-d-old wild-type Col-0, *cle42*, *cle42 EBF1ox* and *EBF1ox*. Bar, 1 cm. (h-j) Chlorophyll content (h), Fv/Fm ratio (i), and *SAG12* expression (j) in the fourth leaves shown in the plants in (g). The bars indicate the mean ± SD (n = 20 for chlorophyll content and Fv/Fm, n = 3 for qPCR). Different letters above the bars indicate statistically significant differences (adjusted P < 0.05, one-way ANOVA).

Given that CLE42p promotes the protein accumulation of EBF1 and thus compromises EIN3 protein stability, we next analyzed whether overexpression of *EBF1* could suppress the premature senescence phenotype of *cle42* mutant. To this end, we examined the leaf senescence phenotypes of Col-0, *cle42*, *cle42 EBF1ox*, and *EBF1ox* plants. Compared with Col-0 and *cle42* mutant, *EBF1ox* plants showed delayed senescence phenotypes with higher chlorophyll content and Fv/Fm, as well as lower levels of *SAG12* gene expression (Fig. 5g–j), indicating that EBF1 delays leaf senescence. More interestingly, overexpression of *EBF1* also suppressed the early senescence phenotypes of *cle42* mutant (Fig. 5g–j), suggesting that EBF1 acts downstream of CLE42 in the regulation of leaf senescence. Because CLE42 promotes EBF1 protein accumulation by suppressing ethylene biosynthesis, we examined whether loss of function of ACSs could rescue the early senescence phenotypes of *cle42*. Toward this end, the *ACS octuple* mutant (*acs2-1 acs4-1 acs5-2 acs6-1 acs7-1 acs9-1 amiRacs8 acs11*) (Tsuchisaka et al., 2009), eliminating the functional redundancy among multiple *ACS* genes, was used to generate *octuple cle42* mutant using the CRISPR-Cas9 technology. As expectedly, mutation of multiple ACSs evidently suppressed the earlier senescence phenotypes of *cle42* mutant (Fig. S14). Together, our results reveal that CLE42 delays leaf senescence through controlling the ACS-EBF-EIN3 cascade.

## Discussion

Leaf senescence is characterized by the transition from nutrient assimilation to remobilization and has a critical impact on agriculture (Gepstein, 2004; Avila-Ospina *et al.*, 2014). As the last stage of leaf development, leaf senescence is not a passive and deteriorative process but a highly regulated and active process (Gepstein, 2004; Lim *et al.*, 2007). The dramatic shift from a photosynthetically active organ to a senescent leaf is induced by a series of endogenous factors, such as age and plant hormones, as well as environmental cues, such as biotic and abiotic stresses (Gepstein, 2004; Guo & Gan, 2005; Lim *et al.*, 2007; Zhang *et al.*, 2021a). Here, we found that CLE42, a peptide hormone of the CLE family in Arabidopsis (Cock & McCormick, 2001), functions as a negative regulator of leaf senescence through integration with ethylene signaling (Fig. 1 and Fig. 4; Fig. S7), an endogenous modulator of plant aging (Abeles *et al.*, 1988).

Peptide hormones participate in cell-to-cell communication to regulate plant growth and development in response to internal and external cues (Matsubayashi & Sakagami, 2006). As the largest group of such peptides, CLE plays crucial roles in regulating meristem activity in shoots and roots as well as in vascular tissues (Cock & McCormick, 2001; Fiers *et al.*, 2005; Ito *et al.*, 2006; Kondo *et al.*, 2006). Recently, CLE14 was found to delay age-dependent and stress-induced leaf senescence by enhancing JUB1-mediated scavenge of reactive oxygen species (ROS) in Arabidopsis (Zhang et al., 2021b). Here, we found that mutation of CLE42 led to premature senescence, whereas overexpression of *CLE42* resulted in prolonged plant longevity (Fig. 1). Exogenous application of chemically synthesized 12-aa peptides derived from CLE42 motifs functionally mimicked the overexpression of *CLE42* and delayed age-dependent and dark-induced leaf senescence (Fig. 3; Fig. S6), supporting previous findings that peptides containing only the C-terminal CLE domain are sufficient to result in CLE activities (Fiers *et al.*, 2005; Ito *et al.*, 2006; Kondo *et al.*, 2006). Expectedly, the earlier senescence phenotypes of *cle42* mutant were also restored by an application of CLE41/44p under dark conditions (Fig. S6), indicative of the existence of function redundancy between CLE42 and CLE41/CLE44 in regulating leaf senescence. Although CLE41 and CLE44 are involved in the regulation of leaf senescence, their contribution seems relatively minor than that of CLE42 (Fig. 2b,c). One of the interpretations would be that the spatiotemporal expression patterns of CLE42 and CLE41/CLE44 are different (Fig. S2) (Yaginuma et al., 2011), which in turn cause different regulatory effects. We cannot exclude the possibility that the delay in leaf senescence mediated by CLE42 and CLE41/CLE44 is through different signaling pathways or possibly controlled by different receptors because they differ by one amino acid in the CLE motif. In supporting this possibility, previous studies find that CLE42 displays a weaker interaction with PXY compared to CLE41/CLE44 (Zhang et al., 2016), and CLE42 possessed partial activity of TDIF (Hirakawa et al., 2008). CLE42 differs from CLE41/CLE44 only in the second position but displayed a different effect on leaf senescence, raising the possibility that the non-conserved residues may allow CLE peptides to be distinguished by their receptors and regulates different developmental process.

Secreted CLE peptides stimulate intracellular signaling through plasma membrane-localized receptors. The binding of a CLE peptide to its receptor triggers various downstream signaling events. CLE42 and CLE41/CLE44 have been shown to bind the LRR-RLK receptor TDR/PXY (Fisher & Turner, 2007; Hirakawa *et al.*, 2008). Similar to the *cle41 cle42 cle44* triple mutant, mutation in *TDR/PXY* also resulted in reduced plant longevity (Fig.2), suggesting that the CLE41/CLE42/CLE44-PXY signaling module functions as a negative regulator of plant aging. Arabidopsis genome contains two PXY-like genes, *PXL1* and *PXL2* (Fisher & Turner, 2007), which function redundantly in regulating vascular tissue development (Etchells *et al.*, 2013). However, no additional contribution of PXL1/PXL2 in leaf senescence was observed when comparing *pxy* and *pxy pxl1 pxl2* (Fig. S5), implicating that PXY is involved in regulating leaf ageing. Compared with *pxy* mutant, the single mutant *cle41*, *cle42* or *cle44* displayed relatively weaker senescence phenotype (Fig. 2); however, the *cle41 cle42 cle44* mutant exhibited stronger senescence phenotype than *pxy* (Fig. 2b–d; Fig. S5), suggesting that the role of CLE41/CLE42/CLE44 in the regulation of leaf senescence might be partially PXY-independent manner. In supporting this hypothesis, overexpression of *CLE42* could slightly rescue the senescence phenotypes of *pxy* (Fig. 3d–f). Previous studies found that CLE9/10 regulate two different processes by being perceived by two distinct receptor systems. CLE9/10 participates in regulating stomatal development through associating with receptor kinase HAESA-LIKE 1 (HSL1), whereas CLE9/10 regulates xylem development through interacting with BARELY NO MERISTEM (BAMs) (Qian et al., 2018). Therefore, more research is needed in the future to explore whether CLE41/42/44 interact with other receptors to control leaf senescence.

CLE peptides regulate a number of biological processes by integrating phytohormone signaling in plants (Wang *et al.*, 2015). CLE10 inhibits protoxylem formation by suppressing the expression of *ARR5* and *ARR6*, two negative regulators of cytokinin signaling (Kondo *et al.*, 2011). Overexpression of *CLE14* or *CLE20* reduces cytokinin content and impacts root growth, which can be rescued by the exogenous application of cytokinin (Meng & Feldman, 2010). Here, we found that CLE42 is involved in multiple factors-induced senescence, including age, darkness, plant hormones (JA and SA) and H_2_O_2_ (Fig.1 and 3; Fig. S3). Interestingly, CLE14 also delays these factors-induced leaf senescence by governing H_2_O_2_ homeostasis (Zhang et al., 2021b). These findings suggest that CLE14 and CLE42 regulate leaf senescence probably through similar pathways. Future studies such as the generation of *cle14 cle42* double mutants need to be conducted to test this possibility. The CLE25 peptide modulates ABA biosynthesis by inducing NCED3 expression in the leaves to regulate stomatal closure under drought conditions (Takahashi *et al.*, 2018). CLE26 affects the primary root protophloem by modulating auxin signaling (Czyzewicz *et al.*, 2015). CLE40 represses cytokinin signaling by suppressing gene expression in signaling and biosynthesis (Pallakies & Simon, 2014). In this study, we found that several hormone signaling pathways, including JA, SA and ethylene, were suppressed in *CLE42-*overexpressing plants through transcriptome analysis (Fig.S7), suggesting that CLE42 delays leaf senescence by suppressing positive modulators of plant aging. Application of the CLE42 peptide reduced the protein accumulation of EIN3 by stabilizing EBF1 and suppressed the function of EIN3 (Fig. 4). Interestingly, expression levels of ethylene biosynthesis gene ACS2 were suppressed by treatment with CLE42p, resulting in a decrease in ethylene production (Fig. S13). Correspondingly, overexpression of *CLE42* evidently suppressed the response to low concentrations of ACC under dark conditions. In contrast, mutation of CLE42 enhanced the response to ethylene (Fig. S10). Moreover, loss of function of EIN3/EIL1, overexpression of *EBF1*, or mutation of *ACS*s suppressed the earlier senescence of the *cle42* mutant (Fig. 5; Fig. S14), suggesting that CLE42 delays leaf senescence by controlling the ACS-EBF-EIN3 cascade. Additionally, overexpression of *CLE42* attenuated the triple response as well as delayed leaf senescence in the presence of ethylene. Conversely, *cle42* mutants showed enhanced responses and early senescence (Fig. S3, S10, and S11). Taken together, our results suggested that CLE42-PXY signaling module controls leaf senescence by communicating with ethylene pathway. It was also noticed that CLE41/42/44, especially CLE42, plays a prominent role in response to low concentrations of ethylene (Fig. S10, S11). Given that plant produces a low level of ethylene during normal and stress conditions, which is sufficient for plant growth, development, and survival (Catala et al., 2014; Johnson and Ecker, 1998; Vong et al., 2019; Yoon and Chen, 2017), our data suggested that CLE42 might be an important player in plant growth and development through interacting with ethylene pathway.

## Acknowledgments

We gratefully thank Dr. Peter Etchells (Durham University, UK) for the *pxy* and *pxy pxl1 pxl2* mutants. This work was supported by the National Natural Science Foundation of China (32170345 and 31970196 to ZL), and the National Key Research and Development Program of China (No. 2019YFA0903904 to H.G.).

## Author contributions

ZL conceived the project and designed the experiments; HG and XX designed some of the experiments; ZY carried out most of the experiments; ST, YG, and CK prepared DNA construct, measured chlorophyll content and Fv/Fm; TI and SS developed CLE CRISPR/Cas9 mutant collection; HLW performed bioinformatics analysis; QY performed WB analysis; ZL and YZ wrote the manuscript with input from all co-authors. All authors agreed on the final version of the manuscript.

## Data availability

RNA-Seq data are available in the Sequence Read Archive (SRA, https://www.ncbi.nlm.nih.gov/sra/) with the accession number PRJNA804857.

## Supporting Information

**Fig. S1** qPCR analysis of *CLE42* expression in the fourth leaves of 3-week-old Col-0 and *CLE42ox* plants.

**Fig. S2** The expression of *CLE41*/*42*/*44* decreases during leaf aging.

**Fig. S3** CLE42 negatively modulates senescence-regulating signal-triggered leaf senescence.

**Fig. S4** Genotyping analysis of *cle41 cle42 cle44* by sequencing.

**Fig. S5** The *cle41 cle42 cle44* shows earlier senescence phenotypes than the receptor mutants.

**Fig. S6** CLE41/44p restored the early senescence phenotype of the *cle42* mutant under dark conditions.

**Fig. S7** Gene ontology analysis of differentially regulated genes in *CLE42ox* plants.

**Fig. S8** The expression of senescence-associated genes in Col-0 and *CLE42ox* plants.

**Fig. S9** Overexpression of *CLE42* caused ethylene insensitivity under low-concentration ACC conditions.

**Fig. S10** Loss of function of CLE42 results in the most prominent triple responses compared with CLE41/CLE44.

**Fig. S11** The expression of *EIN3* in Col-0 upon CLE42p treatment.

**Fig. S12** The expression of downstream target genes of ethylene signaling in Col-0 and *CLE42ox* plants.

**Fig. S13** CLE42 suppresses ethylene biosynthesis.

**Fig. S14** The *ACS octuple* mutant suppressed the earlier senescence phenotypes of *cle42* mutant.

**Table S1.** Primers used in the study.

**Dataset S1.** Differentially expressed genes in *CLE42ox* compared to Col-0.

