## Supporting information for "CLE42 Delays Leaf Senescence by Antagonizing Ethylene Pathway in *Arabidopsis*"

### Supplemental Figures

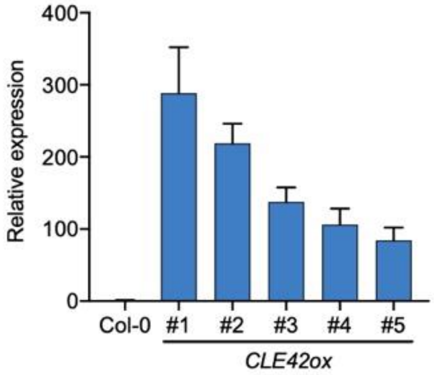

**Fig. S1** qPCR analysis of *CLE42* expression in the fourth leaves of 3-week-old Col-0 and *CLE42ox* plants. Five transgenic lines were used. The bars represent the standard deviation of three technical replicates. Each experiment was repeated at least twice with similar results.

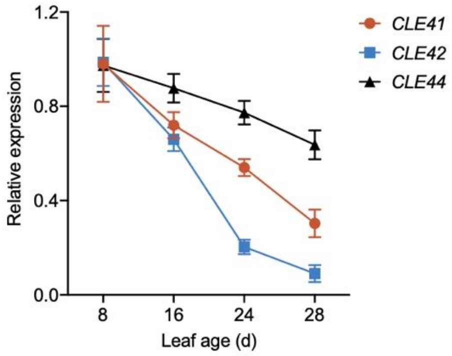

**Fig. S2** The expression of *CLE41*/*42*/*44* decreases during leaf aging. Transcript levels of *CLE41*/*42*/*44* in the fourth leaves of the indicated leaf age. Data are represented as means ± SD (n = 3). **P* < 0.05, ***P* < 0.01, ns, not significant, one-way ANOVA.

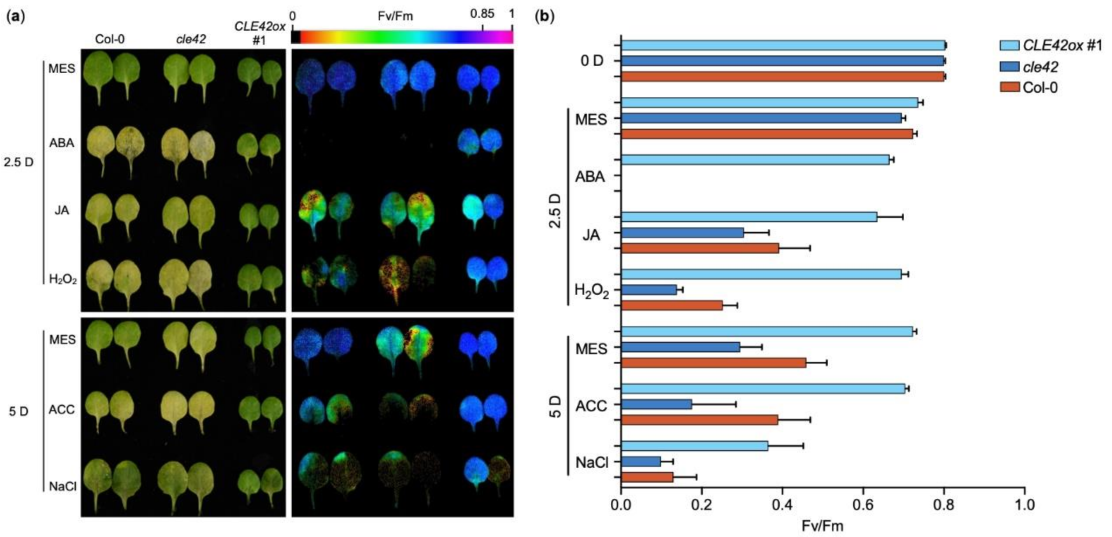

**Fig. S3** CLE42 negatively modulates senescence-regulating signals-triggered leaf senescence. (a) The senescence phenotypes and Fv/Fm image of Col-0, *cle42*, and *CLE42ox* #1 leaves upon treatment with indicated stress-inducing factors under darkness for 2.5 or 5 days. (b) Fv/Fm ratio in leaves shown in (a). The bars indicate mean ± SD (n = 20).

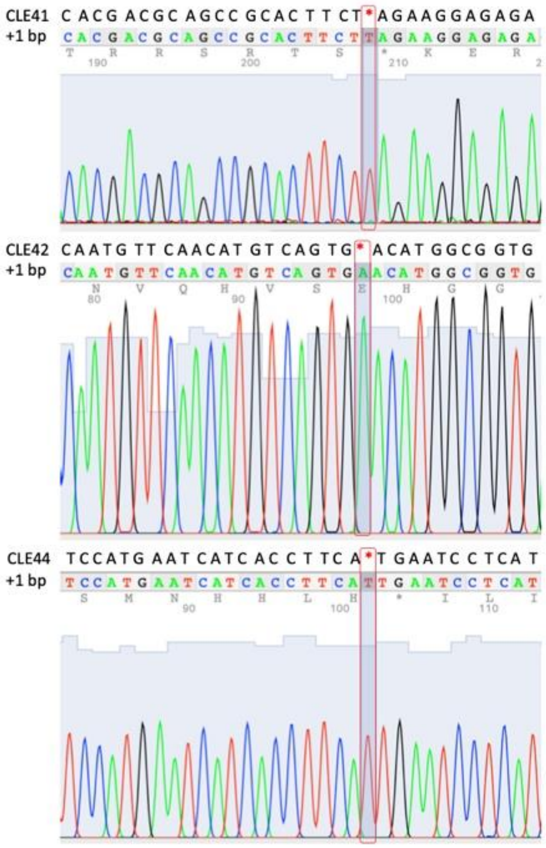

**Fig. S4** Genotyping analysis of *cle41 cle42 cle44* by sequencing. DNA sequencing chromatograms for *CLE41*, *CLE42*, and *CLE44* genes in *cle41 cle42 cle44* triple mutant compared with Col-0. A "T", "A", and "T" insertion is in *cle41*, *cle42*, and *cle44*, respectively.

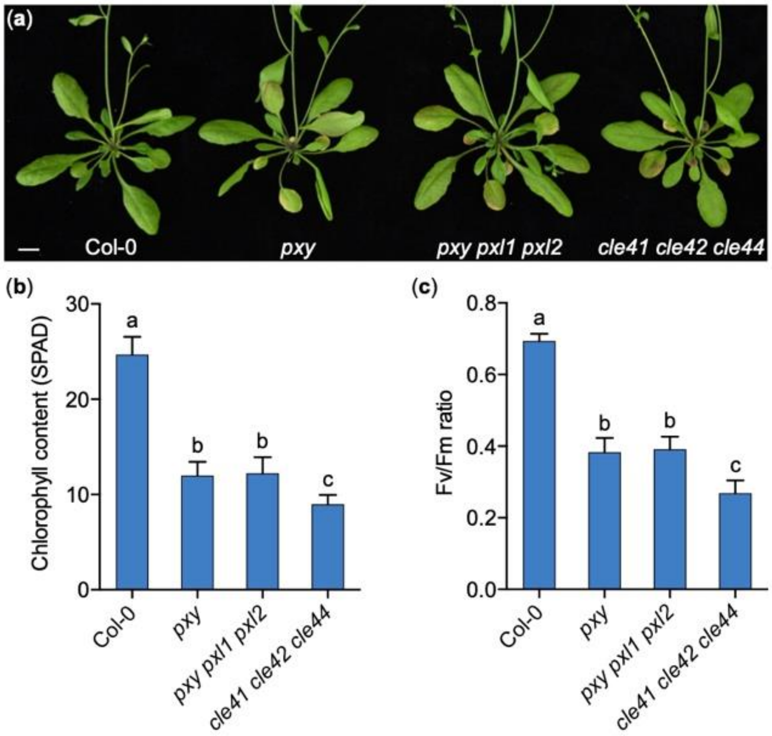

**Fig. S5** The *cle41 cle42 cle44* shows earlier senescence phenotypes than the receptor mutants. (a) The senescence phenotypes of 30-d-old Col-0, *pxy*, *pxy pxl1 pxl2* and *cle41 cle42 cle44*. Bar = 1 cm. (b) and (c) Chlorophyll content (b) and Fv/Fm ratio (c) in the fourth leaves shown in (a). The bars indicate mean ± SD (n = 20). Different letters above the bars indicate statistically significant differences (adjusted $P < 0.05$, one-way ANOVA).

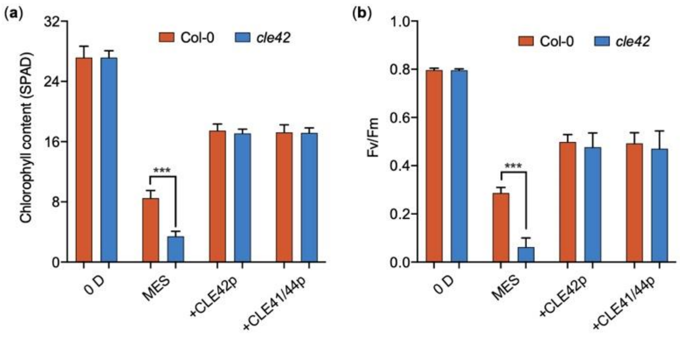

**Fig. S6** CLE41/44p could restore the early senescence phenotype of *cle42* mutant under dark condition. (a) and (b) chlorophyll content (a) and Fv/Fm ratio (b) in the leaves of Col-0 and *cle42* upon CLE42p or CLE41/44p treatment under dark for 5 days. The bars indicate mean ± SD (n = 20). ***P < 0.001, two-way ANOVA.

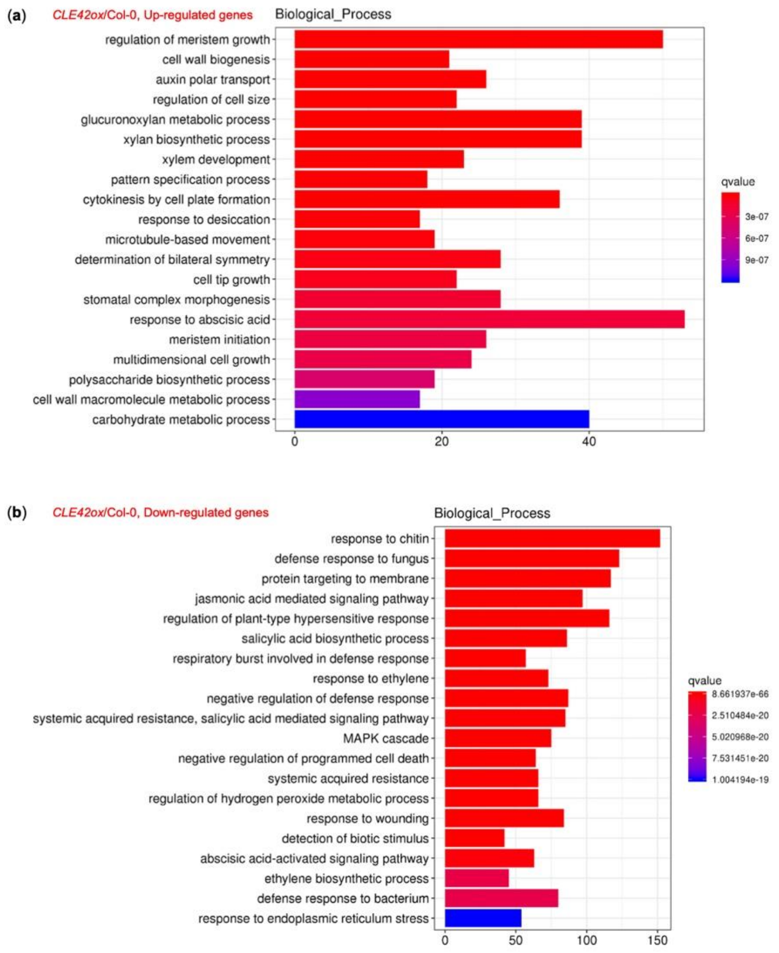

**Fig. S7** Gene ontology analysis of differentially regulated genes in *CLE42ox* plants. (a) and (b) Biological process analyses of the up-regulated (a) and down-regulated (b) genes in *CLE42ox* #1 plants compared with Col-0. Ranking of top 20 biological processes of differentially expressed genes were shown.

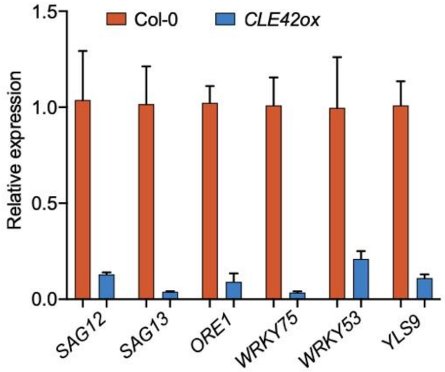

**Fig. S8** The expression of senescence-associated genes in Col-0 and *CLE42ox* plants. Leaves detached from 3-week-old Col-0 and *CLE42ox* #1 plants for qPCR analysis. The bars represent the standard deviation of three technical replicates. Each experiment was repeated at least three times with similar results.

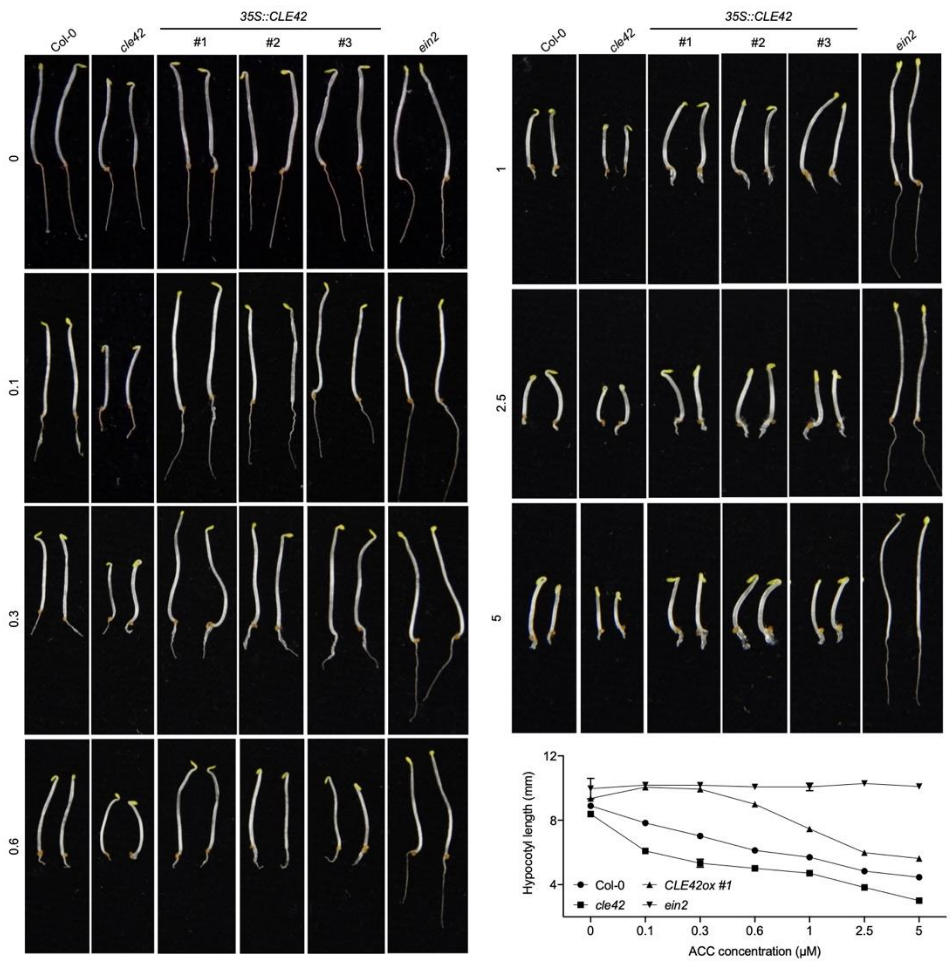

**Fig. S9** Overexpression of *CLE42* caused ethylene insensitive under low concentration of ACC condition. The triple response phenotype of 4-d-old etiolated seedlings (Col-0, *cle42*, and *CLE42ox* #1) grown on MS medium alone or supplemented with different concentration of ACC. Graphical quantification of hypocotyl length in response to various concentrations of ACC. Each bar represents the average length (±SD) of at least 20 seedlings.

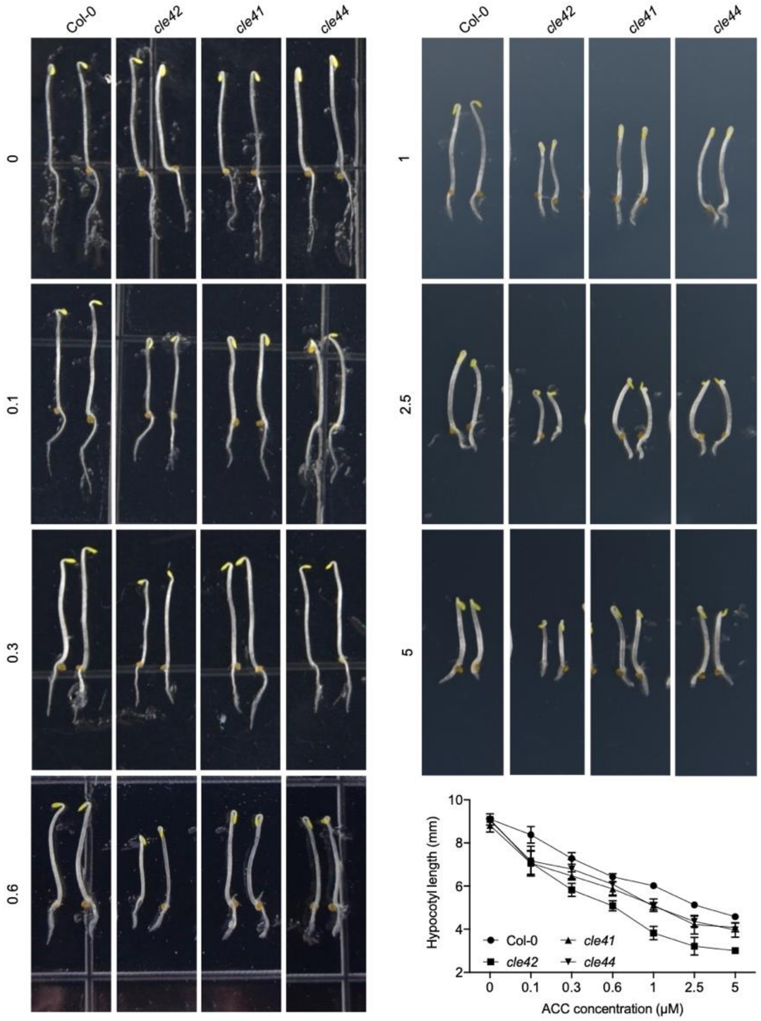

**Fig. S10** Loss function of CLE42 results in most prominent triple responses compared with CLE41/CLE44. The triple response phenotype of 4-d-old etiolated seedlings (Col-0, *cle42*, and *CLE42ox* #1) grown on MS medium alone or supplemented with different concentration of ACC. Graphical quantification of hypocotyl length in response to various concentrations of ACC. Each bar represents the average length (±SD) of at least 20 seedlings.

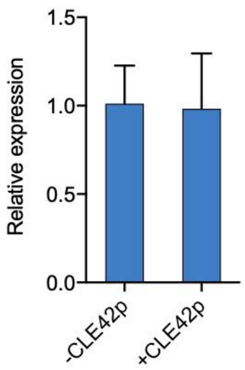

**Fig. S11** The expression of *EIN3* in Col-0 upon CLE42p treatment. 5-d-old seedlings grown on MS with or with 10 μM CLE42p were collected for qPCR analysis. The bars represent the standard deviation of three technical replicates. Each experiment was repeated at least three times with similar results.

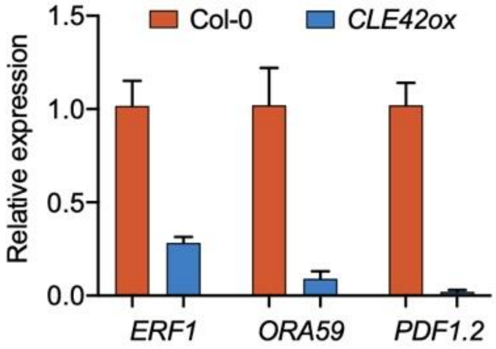

**Fig. S12** The expression of downstream target genes of ethylene signaling in Col-0 and *CLE42ox* plants. Leaves detached from 3-week-old Col-0 and *CLE42ox* #1 plants for qPCR analysis. The bars represent the standard deviation of three technical replicates. Each experiment was repeated at least three times with similar results.

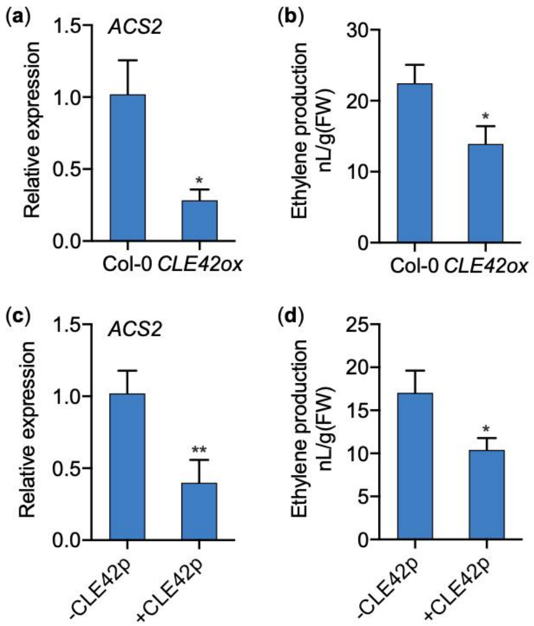

**Fig. S13** CLE42 suppresses ethylene biosynthesis. (a) and (b) qPCR analysis of transcript of *ACS2* (a) and ethylene production (b) in 28-d-old Col-0 and *CLE42ox* #1 plants. The bars indicate mean ± SD (n = 3). (c) and (d) qPCR analysis of transcript of the *ACS2* (c) and ethylene production (d) upon treatment with CLE42p. The third/forth leaves of 28-d-old Col-0 plants were subjected to CLE42p (5 μM, +CLE42p) or water treatment (-CLE42p), and samples were collected at 24 h post incubation for gene expression analysis. *$p < 0.05$, **$p < 0.01$, one-way ANOVA.

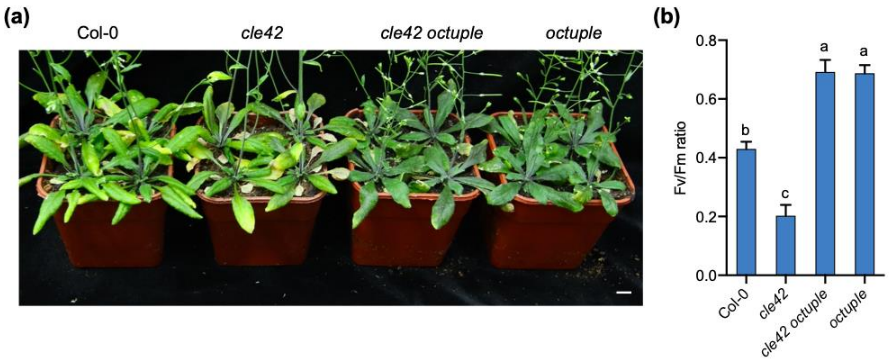

**Fig. S14** The *ACS octuple* mutant suppressed the earlier senescence phenotypes of *cle42* mutant. The senescence phenotypes of 45-d-old wild-type Col-0, *cle42*, *cle42 octuple,* and *octuple* (*acs2-1 acs4-1 acs5-2 acs6-1 acs7-1 acs9-1 amiRacs8 acs11*). Bar, 1 cm. (b) Fv/Fm ratio in the fourth leaves shown in the plants in (a). The bars indicate the mean ± SD (n = 20). Different letters above the bars indicate statistically significant differences (adjusted P < 0.05, one-way ANOVA).

**Table S1.** Primers used in the study.

| Names | Sequence (5'-3') |
|---|---|
| **Plasmid construction** | |
| CLE42OXF | GCGGCCGAATTCCCCGGGATGAGATCTCCTCACATCACCA |
| CLE42OXR | AAGCTTCTCGAGCCCGGGCTACCTATTGGAGATGGGATTT |
| iCLE42F | ATACGCGTTAATTAACTAGTATGAGATCTCCTCACATCACCA |
| iCLE42R | GGAGGCCTGGATCGACTAGTCTACCTATTGGAGATGGGATTT |
| CLE42-CRISPR site | GTTCAACATGTCAGTGACA |
| **Gene expression analysis** | |
| qSAG12F | CAGCTGCGGATGTTGTTG |
| qSAG12R | CCACTTTCTCCCCATTTTG |
| qCLE42F | CCAAACCCATCAAAGAACCATT |
| qCLE42R | CCTATTGGAGATGGGATTTGGA |
| qCLE41F | TCGTCATCAGTCCACATCTATG |
| qCLE41R | GAACCTCATGAGCATCATTTCC |
| qCLE44F | CCATCAAGTCATCCGTTTGATC |
| qCLE44R | GTTGTAGAAGGAACCTCTTGGA |
| qERF1F | TTTCTCTACGGTCTAATCGAGC |
| qERF1R | TGACTTTCTTGAGCTTACGGAT |
| qORE1F | ACAGCTAAGAACGAATGGGTTA |
| qORE1R | CCATTCGGTTAATGTGTGGATC |
| qWRKY75F | CCGTCAAGAACAACAAGTTCCC |
| qWRKY75R | TATGCTCGAAGTTTTCGGTGGA |
| qWRKY53F | AGCCAAAGGATATTCTCGAGAG |
| qWRKY53R | AGTTCCGTAATTGTAAAACGGC |
| qYLS9F | ACTTTAAACGCGGAGAGGATAT |
| qYLS9R | CTTCAAGTCCCCAAGCTTAAAC |
| qORA59F | TCTTCTTCTTCTTCGACGTTGA |
| qORA59R | GCGTCATAACAACACTCTGTTT |
| qPDF1.2F | CTTATCTTCGCTGCTCTTGTTC |
| qPDF1.2R | TGGGAAGACATAGTTGCATGAT |
| qEIN3F | CTGCAGATCACAACAACTTTGA |
| qEIN3R | CATCCATCGTTCCTACTACTCC |

| | |
|---|---|
| qUBC21F | TCAAATGGACCGCTCTTATC |
| qUBC21R | CACAGACTGAAGCGTCCAAG |
| **ChIP-qPCR** | |
| ERF1F | GAGAAAGAAGTTGAAAGCAGAT |
| ERF1R | CATTATCCTATAATCTTAGGAA |
| TUBULIN2F | GAGCCTTACAACGCTACTCTGTCTGT |
| TUBULIN2R | ACACCAGACATAGTAGCAGAAATCA |
